# SARS-CoV-2 Spike amyloid fibrils specifically and selectively accelerates amyloid fibril formation of human prion protein and the amyloid β peptide

**DOI:** 10.1101/2023.09.01.555834

**Authors:** Johan Larsson, Ebba Hellstrand, Per Hammarström, Sofie Nyström

## Abstract

An increasing number of reports suggest an association between COVID-19 infection and initiation or acceleration of neurodegenerative diseases (NDs) including Alzheimer’s disease (AD) and Creutzfeldt-Jakob disease (CJD). Both these diseases and several other NDs are caused by conversion of human proteins into a misfolded, aggregated amyloid fibril state. The fibril formation process is self-perpetuating by seeded conversion from preformed fibril seeds. We recently described a plausible mechanism for amyloid fibril formation of SARS-CoV-2 spike protein. Spike-protein formed amyloid fibrils upon cleavage by neutrophil elastase, abundant in the inflammatory response to COVID-19 infection.

We here provide evidence of significant Spike-amyloid fibril seeded acceleration of amyloid formation of CJD associated human prion protein (HuPrP) using an *in vitro* conversion assay. By seeding the HuPrP conversion assay with other *in vitro* generated disease associated amyloid fibrils we demonstrate that this is not a general effect but a specific feature of spike-amyloid fibrils. We also showed that the amyloid fibril formation of AD associated Aβ1-42 was accelerated by Spike-amyloid fibril seeds. Of seven different 20-amino acid long peptides, Spike532 (_532_NLVKNKCVNFNFNGLTGTGV_551_) was most efficient in seeding HuPrP and Spike601 (_601_GTNTSNQVAVLYQDVNCTEV_620_) was most effective in seeding Aβ1-42, suggesting substrate dependent selectivity of the cross-seeding activity.

Albeit purely *in vitro*, our data suggest that cross-seeding by Spike-amyloid fibrils can be implicated in the increasing number of reports of CJD, AD, and possibly other NDs in the wake of COVID-19.

## Introduction

Amyloids and viruses are each notorious for their detrimental effect on human health. There are also many ways in which cross-talk between these two disease causing entities resulting in increasing risk of harm to the host (1). Several neurodegenerative diseases (NDs) are intimately related to misfolding and amyloid formation of endogenously expressed proteins. However, research has to date failed to describe why some but not others fall victim to these diseases. In a retrospective study of 800 000 individuals from biobanks in Finland and the UK it was evident that infection from some of our most common viruses, *e*.*g*. Influenza and Herpes zoster was connected to an increased risk of some of the most common NDs such as Alzheimer’s disease (AD) and Parkinson’s disease (PD) (2).

The neurological manifestations of COVID-19 both in the acute phase and as long term sequel of SARS-CoV-2 infection have been widely observed and described during the COVID-19 pandemic (3). The dominating explanation for this is neuroinflammation (4) (5) (6) (7). Neuroinflammation is a common denominator of NDs (8) (9) (10) (11). The specific roles of neuroinflammation in prion disease was recently reviewed (12).

NDs are specifically associated with the misfolding and amyloid formation of intracerebral proteins. AD is linked to formation of extracellular amyloid plaques with the Aβ peptide as main constituent and intracellular neurofibrillary tangles composed of Tau protein. Prion diseases are caused by the misfolding and aggregation of the prion protein, PrP, that is abundant on the extracellular surface of all neuronal cells in mammals. Although each distinct disease is associated with its own set of misfolded proteins, more and more evidence is pointing towards the possibility of cross-seeding, that is that amyloids formed of one type of protein can induce amyloid formation of another protein (13) (14) (15).

Several SARS-CoV-2 proteins are known to form amyloid. The proteins translated from viral open reading frame genes (ORFs) are often intrinsically disordered and/or fold only in the context of the viral particle (16). SARS-CoV-2 ORF6 and ORF10 are amyloidogenic(17) and the resulting amyloids exhibit neurotoxic properties on cultured cells (17). Nucleocapsid proteins (NCAPs) are crucial for the assembly of viral particles. These proteins contain low complexity domains that are important for the self-assembly of the virus particle but the complementarity between adjacent polypeptide chains can also promote the formation of amyloid structure (18). The low complexity domain of SARS-CoV-2 NCAP forms amyloid in vitro (19) and this amyloid was recently suggested as a drug candidate for treatment of COVID-19 (19).

The SARS-CoV-2 Spike proteins forms amyloid when cleaved by neutrophil elastase (20). Elastase is abundant in covid induced inflammation (21) (22). Amyloid derived from SARS-CoV-2 spike protein has the potential to hamper fibrinolysis of seeded fibrin and hence might be one explanation for microclot formation in severe and long COVID-19 (20). Data from Brogna and colleagues demonstrate that Spike protein produced in the host as response to mRNA vaccine, as deduced by specific amino acid substitutions, persists in blood samples from 50% of vaccinated individuals for between 67 and 187 days after mRNA vaccination (23). Such prolonged Spike protein exposure has previously been hypothesized to stem from residual virus reservoirs, but evidently this can occur also as consequence of mRNA vaccination.

Other viruses from different families comprise amyloidogenic proteins (1). As an example, several proteins of Influenza A, causing seasonal flu, are known to be amyloidogenic. Recombinant expressed Pb1-F2 protein forms amyloid *in vitro* and in experimentally infected cells (24). Influenza A non-structured protein 1 (NS1) can also from amyloid *in vitro* (25). Influenza A infection has been shown to induce misfolding of PrP (26).

During the past 3 years, several case reports of Creutzfeldt-Jakob disease (CJD) manifestation in parallel with COVID-19 infection or vaccination have been published (27) (28) (29) (30) (31) (32). Recently it was suggested by Stefano et al that the conversion of PrP^C^ to PrP^Sc^ and the subsequent mitochondrial dysfunction should be considered when addressing the etiology of long COVID-19 (33).

Although AD is a very slowly progressing disease, there are already indications that suggest a connection between COVID-19 infection and downstream risk of AD. Brain atrophy was prevalent in patients infected post COVID-19 for early strains (34). Aβ aggregates were found postmortem in brains of young COVID-19 infected patients (35). COVID-19 imposed 1.69 increased risk for new diagnosis of AD within 360 days at age >65 years (36).

A recent *in vitro* study showed augmented Aβ1-42 fibril formation induced by preformed seeds from SARS-CoV-2 Spike-protein peptide 1058-1068 (37). We herein follow up the concept of cross-seeding of ND proteins with SARS-CoV-2 Spike amyloid fibrils. Cross-seeding is a testable hypothesis and we addressed it in our well-established *in vitro* seeding assays by cross-seeding of human prion protein and Aβ1-42 peptide with seven different amyloid fibrils from peptides of the SARS-CoV-2 spike protein.

## Results

We aimed to determine if amyloids derived from the SARS-CoV-2 Spike protein could provoke a faster fibril formation of the human prion protein (HuPrP) and the human Aβ peptide respectively. Hence we added preformed fibrils of seven different 20-amino acid long peptides derived from amyloidogenic sequences from SARS-CoV-2 Spike protein (Wuhan strain) (20) (Table 1), from here on called Spike-seeds with the number corresponding to the initial amino acid sequence in the spike protein.

**Table 1.** Spike peptide sequences used for making Spike fibril seeds.

| Spike Peptide* | Amino acid sequence** |
| --- | --- |
| Spike192 | FVFKNIDGYFKIYSKHTPIN |
| Spike258 | WTAGAAAYYVGYLQPRTFLLK |
| Spike365 | KKKGGGYSVLYNSASFSTFK |
| Spike532 | NLVKNKCVNFNFNGLTGTGV |
| Spike601 | GTNTSNQVAVLYQDVNCTEV |
| Spike685 | KKKRSVASQSIIAYTMSLGA |
| Spike1166 | LGDISGINASVVNIQKEIDR |

\*Peptides were named after the initial amino acid residue of the 1273 residue SARS-CoV-2 spike protein (Wuhan strain) (20). \*\*Highlighted in gray are non-native amino acids introduced for solubility

### Seeding of HuPrP

Human PrP was prepared from *E. Coli* as previously described (38). The protein was diluted to 5 μM which corresponds to 0.1 mg/ml. The seeding fibrils were added to reach 10% (0.01 mg/ml) (Fig 1A) and 1% (0.001 mg/ml) (Fig 1B). To determine specificity of the seeding effect, control seeds from several amyloidogenic protein (insulin, lysozyme, TTR, Aβ1-42) were added to reach a seed concentration of 10% (0.01mg/ml). Fibrillation was monitored by measuring increase in ThT fluorescence over time. The Spike peptides were dissolved in hexafluoro-isopropanol (HFIP), a solvent that affects the surface tension and therefore might disrupt the air-water interface predicted to be important in the fibrillation of HuPrP under native conditions. Hence HFIP was added to the unseeded reactions in corresponding concentrations.

**Figure 1:**
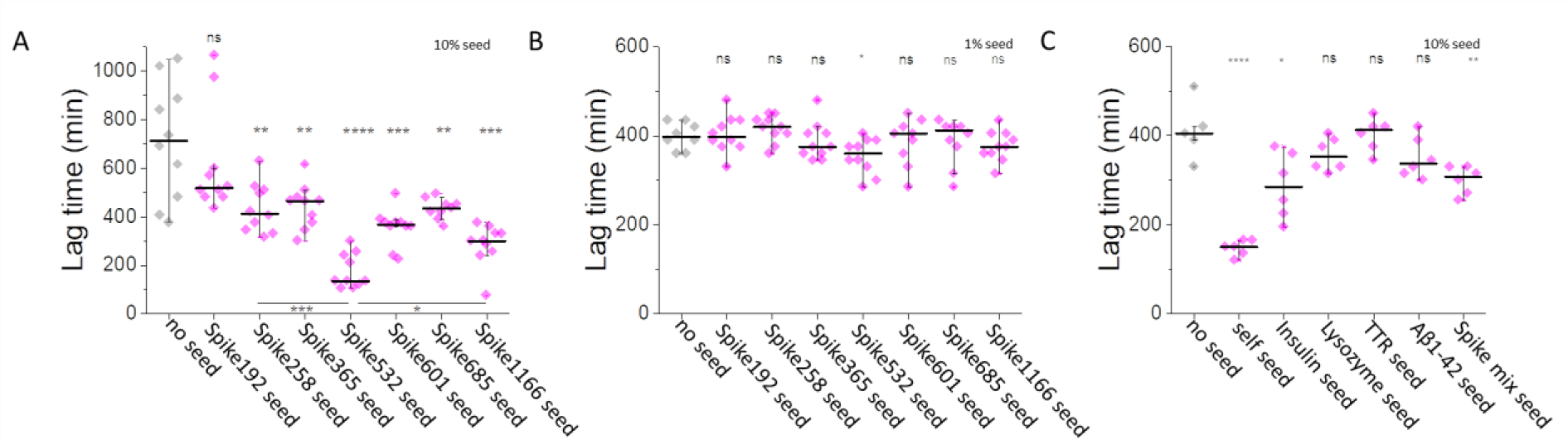
*In vitro* seeding of HuPrP90-231 under native conditions in the presence of Spike amyloid and control seeds. 5 μM (corresponding to ∼0.1 mg/ml) HuPrP was subjected to A) 0.01mg/ml (10 % w/w) Spike fibril amyloid in presence of 1% HFIP as required for the preparation of seeds. Equivalent amount of HFIP was added to the unseeded reaction. B) 0.001 mg/ml (1% w/w) Spike fibrils in presence of 0.1% HFIP as required to prepare the seeds. Equivalent amount of HFIP was added to the unseeded reaction C) 0.01 mg/ml (10% w/w) control seeds from *in vitro* preparations of insulin, lysozyme TTR and Aβ1-42. 0.001 mg/ml (1% w/w) HuPrP90-231 from a previous reaction (self seed) and 0.01mg/ml co-aggregated (1.4 % w/w per constituent peptide) Spike mix were added as positive controls. No HFIP was added to the unseeded reaction. Each reaction was performed in 10 replicates to address the variability in unseeded fibrillation kinetics. The reaction was monitored by ThT fluorescence every 15 minutes during continuous shaking. Lag time was determined as the time point when the ThT intensity had passed 10% of the maximum intensity for that trace.

Addition of 10% preformed fibrils of the seven spike peptides increased the speed of fibrillation for all but one (Fig 1A). Spike192 did not increase the rate of HuPrP fibrillation. We also observed a significant difference in seeding efficacy between the different spike peptide fibrils (Fig 1A).

The HFIP, as predicted, decelerated the fibrillation rate of the unseeded HuPrP reaction at the highest concentration but not at the lower (*c*.*f*. Fig 1 A, B, C unseeded reactions).

A 10-fold dilution of the spike seeds almost totally abolished the seeding potency of the spike peptide amyloids (Fig 1B). However, Spike532 seed still maintained the capacity to significantly decrease the lag time (Fig. 1B). Control reactions with several well-studied amyloids were used to examine the general propensity of cross-seeding HuPrP (Fig. 1C). The control seeds were added to HuPrP to a final concentration of 0,01 mg/ml in correspondence with the highest concentration of spike peptide seeds. 0,001 mg/ml fibrillar HuPrP90-231 was also included as positive control (self seed). Insulin fibrils decreased the lag time of HuPrP fibrillation while fibrils of lysozyme, TTR and Aβ1-42 did not. Spike mix seed comprising a co-fibrillated 14% (1/7th) of each of the seven spike peptides was also included. The spike mix seed (total concentration of 0.01 mg/ml) resulted in a small but significant decrease of HuPrP fibrillation lag time (Fig. 1C).

### Seeding of Aβ1-42

Aβ1-42 was used as a substrate to determine the seeding effect of spike peptide amyloids. The Aβ1-42 concentration was 5 μM (0.022 mg/ml) and the spike peptide amyloids as well as control seeds were added to reach a final concentration of 0.005 mg/ml. We found that all seven spike peptide amyloids efficiently accelerated formation of Aβ1-42 fibrils (Fig 2A). As was the case when seeding HuPrP there is a significant difference between the seeding efficacy between the different spike peptide amyloids. However, Aβ1-42 was more sensitive to Spike601. To ascertain that the seeding effect was not a general effect based on a general amyloid structure we subjected Aβ1-42 to a panel of control seeds and found no acceleration of fibril formation for either insulin, lysozyme or TTR (Fig 2B). Aβ1-42 fibrils and a mix of all seven spike peptides fibrillated together were used as positive controls and both showed seeding activity.

**Figure 2:**
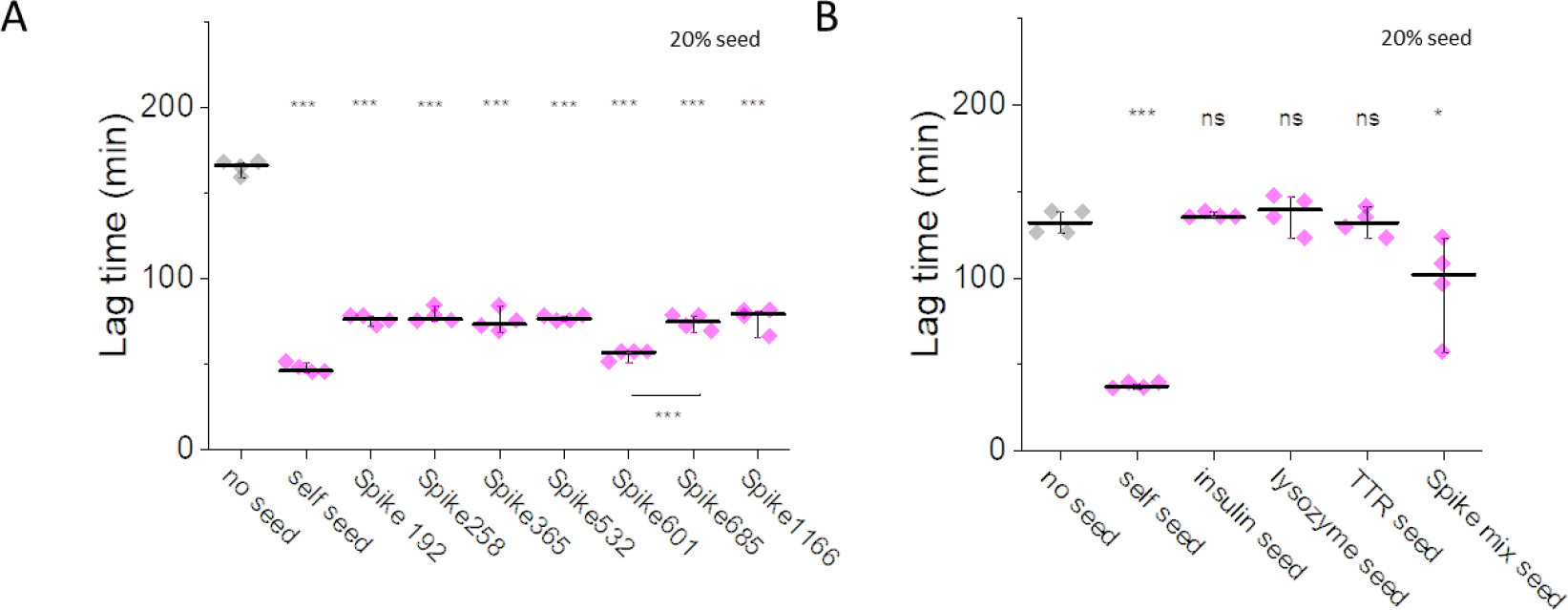
*In vitro* seeding of Aβ1-42 under native conditions in the presence of Spike amyloid and control seeds. 5 μM (corresponding to ∼0.022 mg/ml Aβ1-42) was subjected to A) 0.005mg/ml (20 % w/w) Spike fibril amyloid in presence of 1% HFIP as required for the preparation of seeds. Equivalent amount of HFIP was added to the unseeded reaction. B) 0.01 mg/ml (20% w/w) control seeds from *in vitro* preparations of insulin, lysozyme and TTR. 0.0022 mg/ml (1% w/w) Aβ1-42 from a previous fibrillation reaction (self seed) and 0.01 mg/ml co-aggregated (1.4 % w/w per constituent peptide) Spike mix were added as positive controls. No HFIP was added to the unseeded reaction. Each reaction was performed in 4 replicates. The reaction was monitored by ThT fluorescence every third minute with shaking 1 minute between each measurement. Lag time was determined as the time point when the ThT intensity had passed 10% of the maximum intensity for that trace.

### TEM of selected timepoint in Aβ1-42 fibrillation

In parallel to monitoring the Aβ1-42 fibril formation by ThT, samples from replicate reactions were collected for examination using negative stain transmission electron microscopy (TEM). At timepoint 120 min the exponential growth phase was completed for all the seeded samples while the unseeded sample was still in the initial nucleation phase (Fig 3A). Representative TEM micrographs for each sample are shown in Fig 3B. The unseeded sample contained no fibrillar component but abundant oligomeric structures (Fig. 3B). All samples that were seeded with spike peptide amyloids contained amyloid fibrils (Fig. 3B).

**Figure 3:**
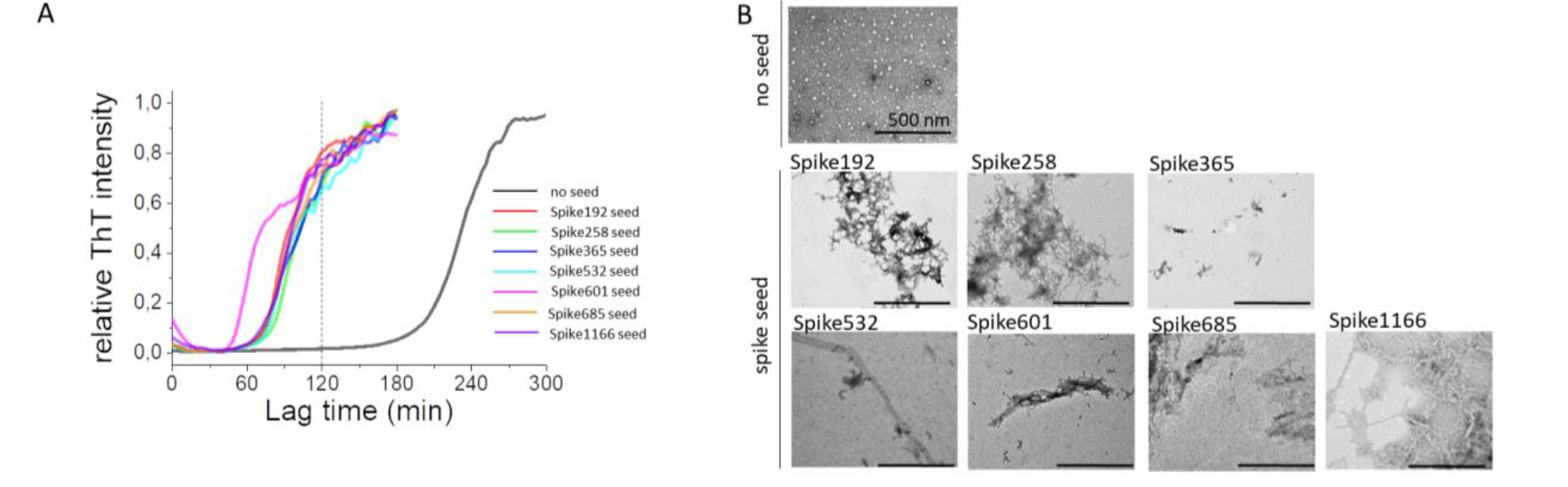
Transmission electron microscopy of Aβ1-42 during *in vitro* seeding by Spike amyloid under native conditions. 5 μM (corresponding to ∼0.022 mg/ml Aβ1-42) was subjected to 0.005mg/ml (20 % w/w) Spike fibril amyloid in presence of 1% HFIP as required for the preparation of seeds. A) Average of 4 kinetic traces for unseeded reaction and each of the Spike amyloid seeded reactions. The grey dotted line at 120 minutes elucidates the timepoint where all the seeded samples are entering the plateau phase of amyloid transition while the unseeded reaction (black) is still not reporting amyloid content by ThT fluorescence. B) shows representative TEM images from all reactions at time point 120 minutes. The Aβ1-42 fibrillation reactions were seeded with the Spike amyloid indicated above each image. All scale bars are 500 nm.

## Discussion

The amyloid fibril formation reactions of recombinant proteins is used as an important research tool in the strive to understand many amyloid-associated diseases. The reaction where a protein is insulted by a small amount of preformed amyloid fibrils of the same protein sequence is known as homologous seeding or self-seeding. The concept of homologous seeding of recombinantly produced and purified proteins is well established and the literature is vast. Seeding assays using recombinant prion protein as substrate are used in clinical diagnostics of prion disease and is known as RT-QuiC (39). If the added amyloid fibrils instead are composed of a different fibril this is called heterologous seeding or cross-seeding. The heterologous seed may be a protein with a single point mutation differing from the substrate, the same protein but from a different species, or a totally different protein but with bearing on the same disease (40). Cross-seeding is rarely as efficient as self-seeding. In many instances, addition of heterologous seeds does not significantly increase the fibril formation of the Aβ peptide (41) (42) (42) or the human prion protein (42) (43). The concept of cross-seeding is currently being explored by many labs and for different disease associated proteins and amino acid complementarity as well as beneficial tertiary (three dimensional) arrangement of the amyloid back bone have been suggested as modulators of success in heterologous seeding (44).

We here demonstrate that both the AD associated peptide Aβ1-42 and the CJD associated human prion protein are susceptible to cross-seeding from amyloids derived from the SARS-CoV-2 Spike protein. For HuPrP the Spike532 amyloid shortens the lag time of amyloid formation significantly more than the other six peptide amyloids and notably Spike192 amyloid did not induce any significant decrease in lag time. The dramatic effect of Spike532 seeding was dose dependent. (All Spike peptide sequences are found in Table 1)

Addition of 10% (0.01 mg/ml in 0.1 mg/ml substrate protein) Spike532 shortened the lag time by almost 80% from a median of 712 minutes to a median of 135 minutes (Fig 1A). A 10-fold dilution of the spike seeds almost totally abolished the seeding activity of the spike peptide amyloids (Fig 1B). However, Spike532 amyloid still maintains the capacity to significantly decrease the lag time but the 10-fold dilution decreased the lag time by only 10% from a median of 397 minutes to a median of 360 minutes. 0.01 mg/ml spike mix, corresponding to 0.0014 mg/ml of the respective peptide decreased the lag time by 25%, from 405 minutes to 307 minutes (Fig 1C).

Control reactions with several well-known and well-studied amyloids were used to examine the general propensity of cross-seeding human prion protein (Fig 1C). The control seeds were added to HuPrP to a final concentration of 0.01 mg/ml in correspondence with the highest concentration of spike peptide amyloids. 0.001 mg/ml fibrillar HuPrP90-231 was included as positive control. Spike mix, comprising 14% (1/7^th^) of each of the seven spike peptides co-aggregated, also at this very low separate peptide concentration did show seeding activity. Insulin fibrils moderately decreased the lag time of HuPrP fibrillation, while fibrils of lysozyme, TTR, and Aβ1-42 did not.

We selected the Aβ1-42 peptide for our experiments as it is most interlinked with AD pathophysiology. Monitoring Aβ as disease biomarker in cerebrospinal fluid it is evident that Aβ1-42/40 ratio in CSF decreases with increasing severity of disease (45). Accumulation of generated Aβ1-42 in amyloid plaques in the brain, a molecular sink effect preventing the peptide from leaking into the CSF is considered a feasible explanation for this biomarker observation. The same shift in ratio between Aβ1-42 and Aβ1-40 was found in COVID-19 patients with neurological manifestations in the early days of the SARS-CoV-2 pandemic (46). The authors provided no molecular explanation for the decreased Aβ1-42/40 ratio in the COVID-19 cases, but accumulation of the Aβ1-42, transient or persistent, is one of several possible explanations and more research is needed to understand this observation.

In our experiments we found that, as was the case for cross-seeding of HuPrP, not all seven Spike amyloid peptides imposed the same acceleration of Aβ1-42 fibrillation. We found that, unlike the case for HuPrP, all our seven tested spike peptide amyloids substantially decreased the time of fibril formation, notably an already rapid unseeded reaction. As in the experiment with HuPrP, one Spike peptide amyloid stood out as most active seed. Intriguingly this sequence was different for Aβ1-42 compared to HuPrP. While Spike532 was most efficient in seeding of HuPrP, Spike601 was most efficient in seeding of Aβ1-42.

The human prion protein is notorious for its sensitivity to self-seeding and is to date the only protein that has an undisputed capacity for inter-individual transmission. On the other hand, seeding the HuPrP with amyloids of heterologous sequences has proven difficult (42) (43).

Both the human prion protein and the Aβ peptide are omnipresent in the human brain but correlate to disease (sporadic CJD and AD respectively) only when in the misfolded state. However, science has to date failed to explain why some but not others fall victim to these diseases. The transmissible properties of PrP prions are well established while inter-host transmission of AD pathology is still under debate.

The evolutionary pressure for viral proteins to strive for complete, successful and quantitative protein folding is low as we recently discussed in a review (1). With similar reasoning, the lack of co-evolution between the human and the viral proteomes, have left humans less protected from cross-seeding from viral amyloids than from self-generated amyloids. There is currently no consensus regarding the etiology of the large number of sporadic cases of NDs in general. Recently Jaunmuktane and colleagues highlighted the vast length of incubation time between insult and disease for vascular Aβ amyloidosis putatively derived from AD donors (47). Herein, alleged victims of iatrogenic transmission of Aβ amyloid seeds presented with Aβ amyloidosis as long as 40+ years after receiving a contaminated dura mater graft or being subjected to contaminated neurosurgical instruments. Putting this together with the rigorous exploration by Levine et al (2) regarding lingering increased risk of NDs decades after viral infection advocates for an intensified research activity on the concept of virus amyloid as a potential triggering factor for protein misfolding and amyloid formation in NDs. Several recent reports of CJD manifesting temporally close to SARS-CoV-2 infection or vaccination with SARS-CoV-2 Spike protein derivatives (27,28,30,31,48) also highlights the need to investigate possible links.

## Experimental procedures

### Protein production and purification

#### HuPrP

All experiments involving human prion protein were performed in a BSL3 lab.

Recombinant HuPrP was expressed and purified as previously described (38)-RSET. In short, E Coli BL21/DE3 were transformed with p-RSET-A plasmid that encoded HuPrP90-231 with an N-terminal hexa-His-tag. The cells were cultured in Terrific broth, induced with IPTG at OD∼2 and harvested 4 hours post induction. The pellet was resuspended in lysis buffer containing 10 mM Tris, 100 mM Na_2_HPO_4_ pH 8, 6 M GuHCl and stored at -80 °C until needed. On day of purification, the pellet was thawed on ice, cells were lysed by sonication and the lysate was cleared by centrifugation at 10 000 g 20 min.

NiNTA agarose beads (Qiagen) were added to the lysate, containing denatured protein. After 30 minutes incubation on rotator, the slurry was poured into a column and the flowthrough was discarded. The agarose beads were washed with the lysis buffer and the protein was refolded by adding gradient mixture of lysis buffer with and without GuHCl. Finally the protein was eluted with 10 mM Tris–HCl, 100 mM Na_2_HPO_4_, 500 mM imidazole, pH 5.8. A second step of purification by size exclusion chromatography on a superdex75 16/60 column was performed to ensure that only natively folded monomeric HuPrP was used for the subsequent fibrillation experiments. Buffer F (50 mM Na_2_HPO_4_ (pH 7.4), 100 mM NaCl, 50 mM KCl) was used as running buffer.

### Aβ1-42

Aβ peptides were purchased from rPeptide. The peptides, lyophilized from HFIP by the vendor, were resuspended in 6 M GuHCL and subjected to size exclusion chromatography using a Superdex 75 10/300 GL. PBS (0.14 M NaCl, 2.7 mM KCl, 0.01 M phosphate and 0.02% (w/v) NaN_3_ pH 7.4) was used as running buffer.

#### Preparation of amyloid seeds

##### Spike peptide seeds

The spike peptides (Table 1) were custom ordered and synthesized by Genscript, Netherlands as previously described in Nyström et al 2022 (20). Lyophilized peptide powder was dissolved in 100% hexafluoroisopropanol (HFIP) to an initial peptide concentration of 10 mg/ml. The peptides, in total 20 amino acids in length, were further diluted in sterile PBS to 1 mg/ml and 10% HFIP. A “spike mix” of all seven peptides was prepared by mixing equal volumes of each peptide of the 1 mg/ml concentration. Samples of each peptide and the peptide mix were incubated at 37 °C in sealed 2 ml non-treated plastic cryotubes (Sarstedt) for 24 hours and amyloid content was determined by ThT fluorescence and TEM (data not shown).

##### TTR fibril seeds

Recombinant native tetrameric TTR-WT protein was expressed and purified as described (49). For concentration determination of tetrameric TTR we used a molar extinction coefficient of 73156 M^-1^cm^-1^ at 280 nm, and a molecular weight of 55570 g/mol .The TTR was dialyzed against milli-Q water at 4 °C using a dialysis tube (Spectra/Por®, Spectrum Laboratories) with a molecular weight cut-off (MWCO) 12–14 kDa and concentrated to 5 mg/ml using spin-filters (Centriprep, Millipore) with a MWCO of 10 kD, at 4 °C. Fibril formation of TTR-WT was performed as described previously (Groenning et al. 2015, PMID: 26108284). Fibril formation was induced by adding a stock solution of pre-cooled (4 °C) 1 M acetic acid, 2 M NaCl to a final concentration of 50 mM acetic acid and 100 mM NaCl (final pH 3.0), to 5 mg/ml TTR in milli-Q water. Samples in sealable 2 mL tubes (Nunc) were mixed briefly and was placed in at 4 °C for several months of stagnant incubation. Formed fibrils were verified by ThT fluorescence and Congo red birefringence under polarized microscopy. TTR Fibril seeds were diluted to 0.2 mg/ml with PBS buffer and were sonicated in water bath prior to seeding experiments.

##### Lysozyme fibril seeds

Fibril seeds of lysozyme was produced according to Mishra et al. 2007 (50). Lyophilized powder of hen egg white lysozyme (Sigma, L6876) was dissolved in 25 mM HCl (pH 1.6) and was dialyzed against 25 mM HCl (pH 1.6) at 4 °C using a porous membrane MWCO 3.5kD (Spectra/Por®, Spectrum Laboratories). The dialyzed protein solution was filtered (0.45 μm pore size filter, Millipore) and protein concentration was determined by absorbance measurements at 280 nm using extinction coefficient ε = 37752 M^-1^cm^-1^, molecular weight of 14300 g/mol. Solutions were prepared in sealable 2 mL tubes (Nunc) by dilution of protein with 25 mM HCl to a final protein concentration of 2.86 mg/ml. Fibril formation of lysozyme was induced by incubation at 65 °C for one week. Formed fibrils were verified by ThT fluorescence and Congo red birefringence under polarized microscopy. Lysozyme Fibril seeds were diluted to 0.2 mg/ml with PBS buffer and were sonicated in water bath prior to seeding experiments.

##### Insulin fibril seeds

Fibril seeds of insulin was produced according to Nilsson et al. 2005 (51). Lyophilized powder of bovine insulin (Sigma, I6634) was dissolved in 25 mM HCl (pH 1.6) and was dialyzed against 25 mM HCl (pH 1.6) at 4 °C using a porous membrane MWCO 3.5kD (Spectra/Por®, Spectrum Laboratories). The dialyzed protein solution was filtered (0.45 μm pore size filter, Millipore) and protein concentration was determined by absorbance measurements at 277 nm using extinction coefficient ε = 5840 M^-1^cm^-1^, molecular weight of 5780 g/mol. Solutions were prepared in sealable 2 mL tubes (Nunc) by dilution of protein with 25 mM HCl to a final protein concentration of 1.85 mg/ml. Fibril formation of insulin was induced by incubation at 65 °C for 24 h. Formed fibrils were verified by ThT fluorescence and Congo red birefringence under polarized microscopy. Insulin fibril seeds were diluted to 0.2 mg/ml with PBS buffer and were sonicated in water bath prior to seeding experiments.

##### Aβ1-42 fibril seeds

Recombinant human Aβ1-42 peptides were purchased from rPeptide (Watkinsville, GA, USA), and were prepared using the same method as described above. The concentration of Aβ1-42 was set to 5 µM and the sample was incubated at 37 °C overnight in quiescent conditions.

#### Cross-seeding experiments

##### Fibrillation of HuPrP

The protein concentration was determined by Abs[280 nm] using extinction coefficient *ε* = 27,515 M^−1^ cm^−1^. The protein was diluted to 6 μM in Buffer F and supplemented with ThT to a final concentration of 2 μM. Aliquotes sufficient for 6 replicates were prepared for each experiment and Spike seeds or control seeds were added to reach the desired concentration for each experiment. The replicate samples were distributed in a non-treated black 96-well plate with clear botton (Corning costar 3880) and sealed firmly with an aluminum foil and a transparent vinyl film to avoid evaporation of HFIP during the experiment.

The fibrillation was performed and monitored in a Tecan Saphire 2 plate reader. The plate was shaken linearly with high intensity for 13 min and then rest for 2 min while measuring. For measurement the samples were excited at 440 nm and fluorescence emission intensity was measured at 480 nm. The lag time of each experiment was defined as the first time point where the fluorescence intensity reached over 10 % of the maximum intensity of that sample. Two-sample T-test was used to evaluate the significance of sample differences.

##### Fibrillation of Aβ1-42

The concentration of Aβ1-42 was determined using the extinction coefficient *ε =* 1,490 M^−1^ cm^−1^ (Abs[275 nm]-Abs[300 nm]) /1,490 M^−1^ cm^−1^. Aβ1-42 was diluted to 5 µM and ThT was added to a final concentration of 2 µM. Spike seeds and control seeds were sonicated for 20 minutes using water bath sonication and then added to the appropriate concentration to each sample. A Tecan Infinite M1000 Pro plate reader was used to monitor the ThT kinetics. excitation at 440 nm and emission at 480 nm was used to monitor the ThT kinetics. Measurements were conducted every third minute and shaking was conducted for 1 minute every third minute. All ThT kinetics were done in 37 °C in a Corning Costar 3880, 96 well plate. The lag time of each experiment was defined as the first time point where the fluorescence intensity reached over 10 % of the maximum intensity of that sample. Two-sample T-test was used to evaluate the significance of sample differences.

#### Transmission Electron Microscopy

Amyloid fibril content at 2 hours was examined by Transmission Electron Microscopy (TEM). Fibrillation reactions for TEM were prepared in parallel to the plate reader reactions using non-treated, well-sealed cryotubes (Sarstedt). The tubes were kept under stagnant conditions at 37 °C in an incubator. Samples for TEM were collected every second hour and the grids were prepared immediately to stop the reaction. Carbon-B grids (formvar carbon coated copper 400 mesh) from Ted Pella were used. 5 µl the fibril sample was placed on a grid for 2 minutes and the excess was removed using a filter paper. Remaining salt was removed from the grid by adding 5 µl milliQ-water to the grid and extract with filter paper. Lastly, 5 µl of 2% aqueous uranyl acetate was placed on the grid for 30 seconds and then the excess was removed with filter paper. TEM images were collected using a Jeol JEM1400 Flash TEM operating at 80 kV.

## Acknowledgments

Instrument time was facilitated by MedFak and ProLinC core facilities at Linköping University

## Author contributions

JL and EH designed and performed experiments and assisted in manuscript preparation, PH contributed necessary funds, planned, designed, and performed experiments, interpreted the data, assisted in manuscript preparation, SN conceptualized the project planned, designed, and performed experiments drafted the manuscript.

## Funding and additional information

Funded by the Swedish research council grant number 2019-04405

## Conflict of interest

The authors declare no conflict of interest.

## Notes

### Competing Interest Statement

The authors have declared no competing interest.

## References

1. Hammarstrom, P., and Nystrom, S. (2023) Viruses and amyloids - a vicious liaison. Prion 17, 82–104

2. Levine, K. S., Leonard, H. L., Blauwendraat, C., Iwaki, H., Johnson, N., Bandres-Ciga, S., Ferrucci, L., Faghri, F., Singleton, A. B., and Nalls, M. A. (2023) Virus exposure and neurodegenerative disease risk across national biobanks. Neuron 111, 1086–1093 e1082

3. Monje, M., and Iwasaki, A. (2022) The neurobiology of long COVID. Neuron 110, 3484–3496

4. Vanderheiden, A., and Klein, R. S. (2022) Neuroinflammation and COVID-19. Curr Opin Neurobiol 76, 102608

5. Soung, A. L., Vanderheiden, A., Nordvig, A. S., Sissoko, C. A., Canoll, P., Mariani, M. B., Jiang, X., Bricker, T., Rosoklija, G. B., Arango, V., Underwood, M., Mann, J. J., Dwork, A. J., Goldman, J. E., Boon, A. C. M., Boldrini, M., and Klein, R. S. (2022) COVID-19 induces CNS cytokine expression and loss of hippocampal neurogenesis. Brain 145, 4193–4201

6. Almutairi, M. M., Sivandzade, F., Albekairi, T. H., Alqahtani, F., and Cucullo, L. (2021) Neuroinflammation and Its Impact on the Pathogenesis of COVID-19. Frontiers in Medicine 8

7. Braga, J., Lepra, M., Kish, S. J., Rusjan, P. M., Nasser, Z., Verhoeff, N., Vasdev, N., Bagby, M., Boileau, I., Husain, M. I., Kolla, N., Garcia, A., Chao, T., Mizrahi, R., Faiz, K., Vieira, E. L., and Meyer, J. H. (2023) Neuroinflammation After COVID-19 With Persistent Depressive and Cognitive Symptoms. JAMA Psychiatry 80, 787–795

8. Guzman-Martinez, L., Maccioni, R. B., Andrade, V., Navarrete, L. P., Pastor, M. G., and Ramos-Escobar, N. (2019) Neuroinflammation as a Common Feature of Neurodegenerative Disorders. Frontiers in Pharmacology 10

9. Mohamed, W., Kumar, J., Alghamdi, B. S., Soliman, A.-H., and Toshihide, Y. (2023) Neurodegeneration and inflammation crosstalk: Therapeutic targets and perspectives. IBRO Neuroscience Reports 14, 95–110

10. Kempuraj, D., Thangavel, R., Natteru, P. A., Selvakumar, G. P., Saeed, D., Zahoor, H., Zaheer, S., Iyer, S. S., and Zaheer, A. (2016) Neuroinflammation Induces Neurodegeneration. J Neurol Neurosurg Spine 1

11. Kwon, H. S., and Koh, S.-H. (2020) Neuroinflammation in neurodegenerative disorders: the roles of microglia and astrocytes. Translational Neurodegeneration 9, 42

12. Li, B., Chen, M., and Zhu, C. (2021) Neuroinflammation in Prion Disease. Int J Mol Sci 22

13. Chaudhuri, P., Prajapati, K. P., Anand, B. G., Dubey, K., and Kar, K. (2019) Amyloid cross-seeding raises new dimensions to understanding of amyloidogenesis mechanism. Ageing Res Rev 56, 100937

14. Ren, B., Zhang, Y., Zhang, M., Liu, Y., Zhang, D., Gong, X., Feng, Z., Tang, J., Chang, Y., and Zheng, J. (2019) Fundamentals of cross-seeding of amyloid proteins: an introduction. J Mater Chem B 7, 7267–7282

15. Subedi, S., Sasidharan, S., Nag, N., Saudagar, P., and Tripathi, T. (2022) Amyloid Cross-Seeding: Mechanism, Implication, and Inhibition. Molecules 27

16. Mateu, M. G. (2013) Assembly, stability and dynamics of virus capsids. Arch Biochem Biophys 531, 65–79

17. Charnley, M., Islam, S., Bindra, G. K., Engwirda, J., Ratcliffe, J., Zhou, J., Mezzenga, R., Hulett, M. D., Han, K., Berryman, J. T., and Reynolds, N. P. (2022) Neurotoxic amyloidogenic peptides in the proteome of SARS-COV2: potential implications for neurological symptoms in COVID-19. Nat Commun 13, 3387

18. Langenberg, T., Gallardo, R., van der Kant, R., Louros, N., Michiels, E., Duran-Romaña, R., Houben, B., Cassio, R., Wilkinson, H., Garcia, T., Ulens, C., Van Durme, J., Rousseau, F., and Schymkowitz, J. (2020) Thermodynamic and Evolutionary Coupling between the Native and Amyloid State of Globular Proteins. Cell Rep 31, 107512

19. Tayeb-Fligelman, E., Bowler, J. T., Tai, C. E., Sawaya, M. R., Jiang, Y. X., Garcia, G., Griner, S. L., Cheng, X., Salwinski, L., Lutter, L., Seidler, P. M., Lu, J., Rosenberg, G. M., Hou, K., Abskharon, R., Pan, H., Zee, C.-T., Boyer, D. R., Li, Y., Anderson, D. H., Murray, K. A., Falcon, G., Cascio, D., Saelices, L., Damoiseaux, R., Arumugaswami, V., Guo, F., and Eisenberg, D. S. (2023) Low complexity domains of the nucleocapsid protein of SARS-CoV-2 form amyloid fibrils. Nature Communications 14, 2379

20. Nyström, S., and Hammarström, P. (2022) Amyloidogenesis of SARS-CoV-2 Spike Protein. Journal of the American Chemical Society 144, 8945–8950

21. Liao, D., Zhou, F., Luo, L., Xu, M., Wang, H., Xia, J., Gao, Y., Cai, L., Wang, Z., Yin, P., Wang, Y., Tang, L., Deng, J., Mei, H., and Hu, Y. (2020) Haematological characteristics and risk factors in the classification and prognosis evaluation of COVID-19: a retrospective cohort study. Lancet Haematol 7, e671–e678

22. Antonioli, L., Fornai, M., Pellegrini, C., and Blandizzi, C. (2020) NKG2A and COVID-19: another brick in the wall. Cellular & Molecular Immunology 17, 672–674

23. Brogna, C., Cristoni, S., Marino, G., Montano, L., Viduto, V., Fabrowski, M., Lettieri, G., and Piscopo, M. (2023) Detection of recombinant Spike protein in the blood of individuals vaccinated against SARS-CoV-2: Possible molecular mechanisms. Proteomics Clin Appl, e2300048

24. Chevalier, C., Al Bazzal, A., Vidic, J., Fevrier, V., Bourdieu, C., Bouguyon, E., Le Goffic, R., Vautherot, J. F., Bernard, J., Moudjou, M., Noinville, S., Chich, J. F., Da Costa, B., Rezaei, H., and Delmas, B. (2010) PB1-F2 influenza A virus protein adopts a beta-sheet conformation and forms amyloid fibers in membrane environments. J Biol Chem 285, 13233–13243

25. Shaldzhyan, A. A., Zabrodskaya, Y. A., Baranovskaya, I. L., Sergeeva, M. V., Gorshkov, A. N., Savin, II, Shishlyannikov, S. M., Ramsay, E. S., Protasov, A. V., Kukhareva, A. P., and Egorov, V. V. (2021) Old dog, new tricks: Influenza A virus NS1 and in vitro fibrillogenesis. Biochimie 190, 50–56

26. Hara, H., Chida, J., Uchiyama, K., Pasiana, A. D., Takahashi, E., Kido, H., and Sakaguchi, S. (2021) Neurotropic influenza A virus infection causes prion protein misfolding into infectious prions in neuroblastoma cells. Sci Rep 11, 10109

27. Young, M. J., O’Hare, M., Matiello, M., and Schmahmann, J. D. (2020) Creutzfeldt-Jakob disease in a man with COVID-19: SARS-CoV-2-accelerated neurodegeneration? Brain Behav Immun 89, 601–603

28. Tayyebi, G., Malakouti, S. K., Shariati, B., and Kamalzadeh, L. (2022) COVID-19-associated encephalitis or Creutzfeldt-Jakob disease: a case report. Neurodegener Dis Manag 12, 29–34

29. Olivo, S., Furlanis, G., Buoite Stella, A., Fabris, M., Milanic, R., Zanusso, G., and Manganotti, P. (2023) Rapidly evolving Creutzfeldt-Jakob disease in COVID-19: from early status epilepticus to fatal outcome. Acta Neurol Belg 123, 1553–1556

30. Bernardini, A., Gigli, G. L., Janes, F., Pellitteri, G., Ciardi, C., Fabris, M., and Valente, M. (2022) Creutzfeldt-Jakob disease after COVID-19: infection-induced prion protein misfolding? A case report. Prion 16, 78–83

31. Ciolac, D., Racila, R., Duarte, C., Vasilieva, M., Manea, D., Gorincioi, N., Condrea, A., Crivorucica, I., Zota, E., Efremova, D., Crivorucica, V., Ciocanu, M., Movila, A., and Groppa, S. (2021) Clinical and Radiological Deterioration in a Case of Creutzfeldt-Jakob Disease following SARS-CoV-2 Infection: Hints to Accelerated Age-Dependent Neurodegeneration. Biomedicines 9

32. Alloush, T. K., Alloush, A. T., Abdelazeem, Y., Shokri, H. M., Abdulghani, K. O., and Elzoghby, A. (2023) Creutzfeldt-Jakob disease in a post-COVID-19 patient: did SARS-CoV-2 accelerate the neurodegeneration? Egypt J Neurol Psychiatr Neurosurg 59, 69

33. Stefano, G. B., Büttiker, P., Weissenberger, S., Anders, M., Raboch, J., Ptacek, R., and Kream, R. M. (2023) Potential Prion Involvement in Long COVID-19 Neuropathology, Including Behavior. Cellular and Molecular Neurobiology 43, 2621–2626

34. Douaud, G., Lee, S., Alfaro-Almagro, F., Arthofer, C., Wang, C., McCarthy, P., Lange, F., Andersson, J. L. R., Griffanti, L., Duff, E., Jbabdi, S., Taschler, B., Keating, P., Winkler, A. M., Collins, R., Matthews, P. M., Allen, N., Miller, K. L., Nichols, T. E., and Smith, S. M. (2022) SARS-CoV-2 is associated with changes in brain structure in UK Biobank. Nature

35. Priemer, D. S., Rhodes, C. H., Karlovich, E., Perl, D. P., and Goldman, J. E. (2022) Abeta Deposits in the Neocortex of Adult and Infant Hypoxic Brains, Including in Cases of COVID-19. J Neuropathol Exp Neurol 81, 988–995

36. Wang, L., Davis, P. B., Volkow, N. D., Berger, N. A., Kaelber, D. C., and Xu, R. (2022) Association of COVID-19 with New-Onset Alzheimer’s Disease. J Alzheimers Dis 89, 411–414

37. Cao, S., Song, Z., Rong, J., Andrikopoulos, N., Liang, X., Wang, Y., Peng, G., Ding, F., and Ke, P. C. (2023) Spike Protein Fragments Promote Alzheimer’s Amyloidogenesis. ACS Appl Mater Interfaces 15, 40317–40329

38. Sandberg, A., and Nystrom, S. (2018) Purification and Fibrillation of Recombinant Human Amyloid-beta, Prion Protein, and Tau Under Native Conditions. Methods Mol Biol 1779, 147–166

39. Peden, A. H., McGuire, L. I., Appleford, N. E. J., Mallinson, G., Wilham, J. M., Orru, C. D., Caughey, B., Ironside, J. W., Knight, R. S., Will, R. G., Green, A. J. E., and Head, M. W. (2012) Sensitive and specific detection of sporadic Creutzfeldt-Jakob disease brain prion protein using real-time quaking-induced conversion. J Gen Virol 93, 438–449

40. Mandal, P. K., Pettegrew, J. W., Masliah, E., Hamilton, R. L., and Mandal, R. (2006) Interaction between Aβ Peptide and α Synuclein: Molecular Mechanisms in Overlapping Pathology of Alzheimer’s and Parkinson’s in Dementia with Lewy Body Disease. Neurochemical Research 31, 1153–1162

41. Liang, R., Tian, Y., and Viles, J. H. (2022) Cross-seeding of WT amyloid-beta with Arctic but not Italian familial mutants accelerates fibril formation in Alzheimer’s disease. J Biol Chem 298, 102071

42. Gallardo, R., Ramakers, M., De Smet, F., Claes, F., Khodaparast, L., Khodaparast, L., Couceiro, J. R., Langenberg, T., Siemons, M., Nyström, S., Young, L. J., Laine, R. F., Young, L., Radaelli, E., Benilova, I., Kumar, M., Staes, A., Desager, M., Beerens, M., Vandervoort, P., Luttun, A., Gevaert, K., Bormans, G., Dewerchin, M., Van Eldere, J., Carmeliet, P., Vande Velde, G., Verfaillie, C., Kaminski, C. F., De Strooper, B., Hammarström, P., Nilsson, K. P., Serpell, L., Schymkowitz, J., and Rousseau, F. (2016) De novo design of a biologically active amyloid. Science 354

43. Nystrom, S., and Hammarstrom, P. (2015) Generic amyloidogenicity of mammalian prion proteins from species susceptible and resistant to prions. Sci Rep 5, 10101

44. Morales, R., Moreno-Gonzalez, I., and Soto, C. (2013) Cross-seeding of misfolded proteins: implications for etiology and pathogenesis of protein misfolding diseases. PLoS Pathog 9, e1003537

45. Hansson, O., Lehmann, S., Otto, M., Zetterberg, H., and Lewczuk, P. (2019) Advantages and disadvantages of the use of the CSF Amyloid β (Aβ) 42/40 ratio in the diagnosis of Alzheimer’s Disease. Alzheimer’s Research & Therapy 11, 34

46. Ziff, O. J., Ashton, N. J., Mehta, P. R., Brown, R., Athauda, D., Heaney, J., Heslegrave, A. J., Benedet, A. L., Blennow, K., Checkley, A. M., Houlihan, C. F., Gauthier, S., Rosa-Neto, P., Fox, N. C., Schott, J. M., Zetterberg, H., Benjamin, L. A., and Paterson, R. W. (2022) Amyloid processing in COVID-19-associated neurological syndromes. Journal of Neurochemistry 161, 146–157

47. Jaunmuktane, Z., Banerjee, G., Paine, S., Parry-Jones, A., Rudge, P., Grieve, J., Toma, A. K., Farmer, S. F., Mead, S., Houlden, H., Werring, D. J., and Brandner, S. (2021) Alzheimer’s disease neuropathological change three decades after iatrogenic amyloid-beta transmission. Acta Neuropathol 142, 211–215

48. Olivo, S., Furlanis, G., Buoite Stella, A., Fabris, M., Milanic, R., Zanusso, G., and Manganotti, P. (2022) Rapidly evolving Creutzfeldt-Jakob disease in COVID-19: from early status epilepticus to fatal outcome. Acta Neurol Belg, 1-4

49. Begum, A., Zhang, J., Derbyshire, D., Wu, X., Konradsson, P., Hammarstrom, P., and von Castelmur, E. (2023) Transthyretin Binding Mode Dichotomy of Fluorescent trans-Stilbene Ligands. ACS chemical neuroscience 14, 820–828

50. Mishra, R., Sorgjerd, K., Nystrom, S., Nordigarden, A., Yu, Y. C., and Hammarstrom, P. (2007) Lysozyme amyloidogenesis is accelerated by specific nicking and fragmentation but decelerated by intact protein binding and conversion. J Mol Biol 366, 1029–1044

51. Nilsson, K. P., Herland, A., Hammarstrom, P., and Inganas, O. (2005) Conjugated polyelectrolytes: conformation-sensitive optical probes for detection of amyloid fibril formation. Biochemistry 44, 3718–3724

